# The mismeasure of (obese) man – body mass index is an inadequate indicator of body condition

**DOI:** 10.1101/149088

**Authors:** Lukáš Kratochvíl, Jaroslav Flegr

## Abstract

The body mass index (BMI), recommended also by the World Health Organization, is currently used as the leading body condition indicator in clinical and epidemiological studies and has become popular among the general public. Here we provide evidence of a systematic bias in BMI, showing that BMI is dependent on body height. As a result, shorter persons have a greater chance of being classified as underweight, while taller persons as overweight, even if they have identical nutritional status. Use of BMI should be thus abandoned in diagnosis as well as in clinical and experimental studies.

## Introduction

With the global epidemic of overweight and obesity in developed countries, a simple and straightforward indicator of individual and population nutritional status is urgently needed. For this purpose, the World Health Organization (WHO; 2010) uses the body mass index (BMI) defined as the ratio of the weight in kilograms to the square of the height in meters (kg/m^2^), with the principal cut-off points of <18.50 for underweight and ≥25.00 for overweight. The body mass index is currently used as the leading body condition indicator in clinical and epidemiological studies and has also become popular among the general public as a tool for body condition monitoring. Nevertheless, it has long been known that the ratios between two body measurements can change with the variation in the body size and consequently can lead to biased conclusions (e.g. Huxley & Teissier 1936). For this reason, the use of the ratios has been largely abandoned in most comparative studies, at least in animal morphometrics or ecology. It is therefore surprising that many ratios such as the BMI, waist-to-hip ratio or digit ratio are still widely used in human studies of great medical importance (for criticism see e.g. Kratochvíl & Flegr 2009; Lolli et al. 2017).

Here we provide evidence of a systematic bias in an original set of BMI data showing that BMI is dependent on body height. Therefore, BMI is not appropriate for any comparison of body condition among people of different heights.

## Material and methods

Data on body height and mass were collected from visitors of a public library in Prague, Czech Republic. BMI was compared between men and women by t-test. Next, we fitted the full-factorial GLM model with BMI as a dependent variable, sex as an independent variable and the square of the height in meters as a covariate, and searched for a significant effect of these factors and their interactions with the covariate. All tests were performed in *Statistica vers*. *6.0* (StatSoft Inc. 2001).

## Results and Discussion

The mean BMI was significantly higher in men, 23.73, than in women, 21.58 (t-test: p < 0.000001; n = 997, 532 women, 465 men). However, when we fitted the full-factorial general linear model with BMI as a dependent variable, sex as an independent variable and the square of the height in meters as a covariate, neither sex nor interaction between sex and the covariate had a significant effect. Therefore, no difference in BMI (and obesity) was observed between men and women when an appropriate statistical analysis had been applied. BMI significantly increased with the square of the height following the formula BMI = 17.32 + 1.72*height^2^ (Fig. 1). According to this relationship, the mean BMI for persons with a height of 150 cm is predicted to be 21.19, while it is as high as 24.20 for those 200 cm in height. The increase in BMI with body height is mainly a consequence of the negative intercept in the relationship between the weight in kilograms and the square of the height in meters. In our sample, the weight = −17.00 + 28.19*height^2^, which means that BMI = 28.19 − 17.00/height^2^. As a result, shorter persons, here usually women, have a greater chance of being classified as underweight, while taller persons have a greater chance of being classified as overweight, even if they have identical nutritional status.

**Fig. 1.**
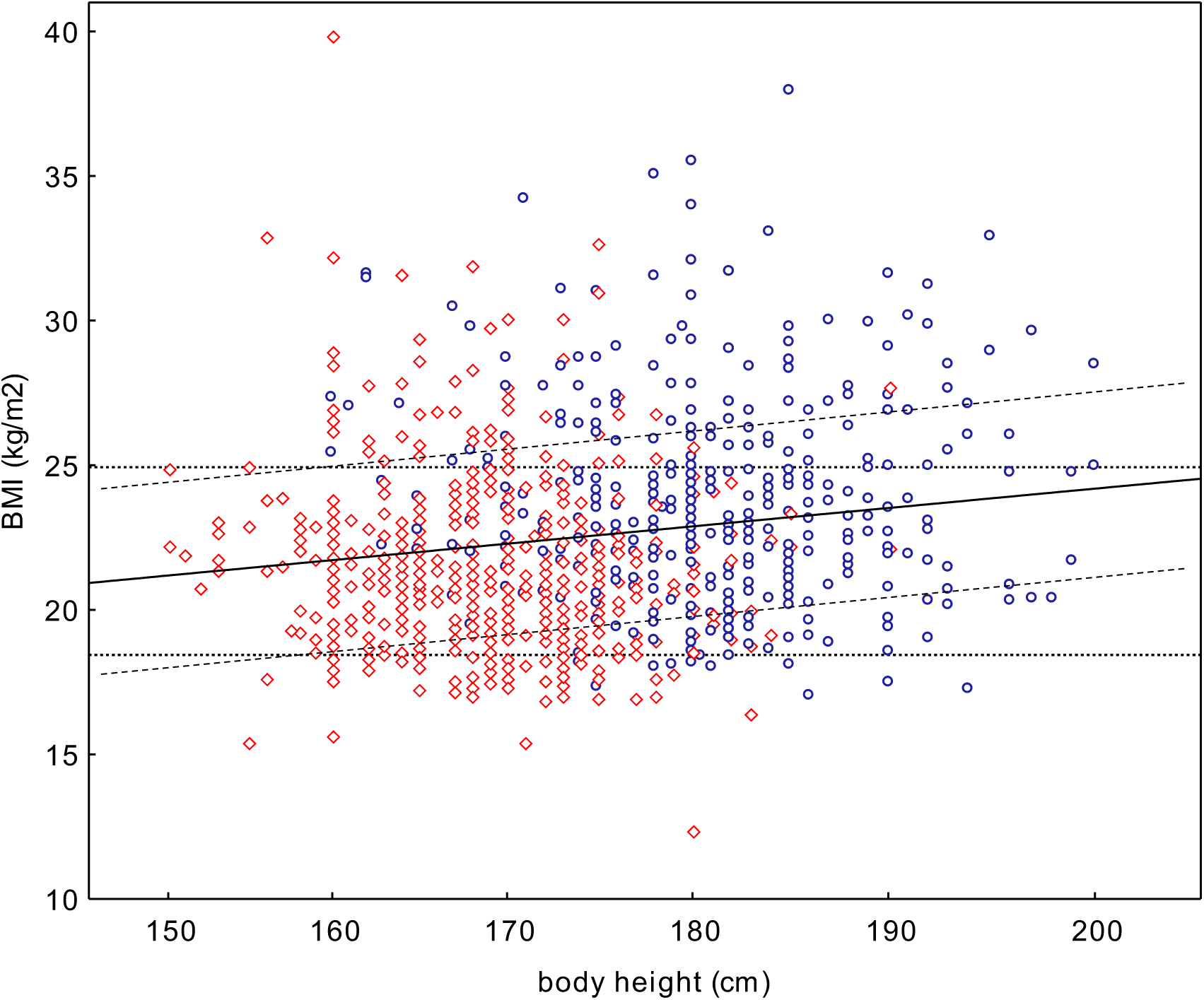
BMI systematically increases with body height. Therefore, the static criterion of two principal BMI cut-off points (18.50 and 25.00; dotted lines) recommended by WHO to identify or overweight persons is inadequate. The correct criterion should take into account the scaling of BMI with body height (dashed lines). In the present dataset, the nutritional status is misclassified in nearly 11% of subjects: 40 subjects of normal nutritional status are misclassified as overweight and 68 underweight subjects are misclassified as normal-weight. Red squares – women, blue circles – men.

In summary, although initially constructed as a measure of body condition controlled for differences in body height, BMI is highly dependent on body height. We stress that - as many other ratios - the body mass index is an inadequate indicator and its use in diagnosis as well as in clinical and experimental studies should be abandoned. Anthropologists, epidemiologists and physicians can draw inspiration from animal ecologists (recently e.g. Peig & Green 2010) and seek for a more appropriate, unbiased indicator of body condition in humans.

## Acknowledgement

The research was supported by GAČR, project no. 16-24619S and Charles University, UNCE 204004.

